# The organizer as a cooperative of signaling cells for neural induction

**DOI:** 10.1101/2025.03.05.641623

**Authors:** Hui-Chun Lu, Katherine E. Trevers, Tatiana Solovieva, Claire Anderson, Linette Pérez-Campos, Lenka Filipkova, Vlad Arimia, Charlotte Colle, Nidia M. M. De Oliveira, Leslie Dale, Claudio D. Stern

## Abstract

The “organizer”, discovered 100 years ago by Hans Spemann and Hilde Mangold, is a special region of vertebrate embryos at the gastrula stage; it emits signals that can re-direct the fate of neighboring cells to acquire neural plate identity. It is generally imagined as unique population of cells producing one or a few signaling molecules, responsible for neural induction and for patterning the neural plate and the mesoderm. Here we use single cell and tissue transcriptomics to explore the expression of signaling molecules in the node (the amniote organizer). Although all organizer cells express the homeobox gene *Goosecoid*, node cells show a diversity of transcription factor signatures associated with expression of subsets of many signaling molecules, suggesting distinct cell sub-populations. Using a recently described Gene Regulatory Network (GRN) of 175 transcriptional responses to neural induction, we explore the activities of 22 of these signals and find that some of them regulate the expression of components of the GRN that are not responsive to previously described pathways associated with neural induction. These results suggest that rather than a single, static, homogeneous population, the organizer comprises a diverse collective of specialized cells that emit cooperating signals to instruct receiving neighbors to adopt their new identities.

**Significance Statement:** The Spemann-Mangold organizer is an embryonic region that can induce the formation of a fully patterned nervous system from non-neural embryonic cells. Here we show that it is made up of a diversity of cell populations that emit distinct sets of signals, which cooperate to account for the repertoire of molecular responses in receiving cells. Several of these signals had not previously been associated with neural induction or patterning.

## Introduction

During gastrulation in vertebrate embryos, the organizer resides at the dorsal-most part of the blastopore in anamniotes and at the anterior tip of the primitive streak (Hensen’s node) in amniotes(1, 2). A key property unique to the organizer, which led to its discovery some 100 years ago(3, 4) is that upon transplantation to a remote region of a host embryo, it can redirect the fate of adjacent non-neural (prospective epidermis or extraembryonic) ectoderm to a neural plate fate. After decades of unsuccessfully trying to identify the signals from the organizer responsible for this induction, inhibition of BMP emerged as playing a key role(5-11). However, because BMP levels critically regulate other aspects of early embryo patterning both earlier and later in development, few experimental approaches have been able to separate these functions from each other. A further complication is that depending on where it is placed, a graft of the organizer can also recruit cells of the host neural plate to form a separate neural plate, but without changing cell fates. The chick embryo offers the advantage that grafts of Hensen’s node to the area opaca, far from the embryo’s neural plate, can induce a complete neural plate. This assay circumvents the problems above and allows precise time-course studies to be performed, since the time of grafting defines “time-zero”. This system revealed that although BMP inhibition is an absolute requirement for neural induction by the node, it is not sufficient to mimic all of the neural inducing effects of a node transplant(12-15). Subsequent experiments in the chick and other model organisms implicated FGFs and other factors including IGF, Wnt-inhibition, retinoids, Notch and hedgehog in conjunction with BMP inhibition(11, 14, 16-19). However, we still don’t know what specific contributions each of these signals makes to the neural induction process. A recent study has uncovered an intricate network of transcriptional responses over time, describing a cascade of expression of 175 transcriptional regulators and predicting more than 5,000 regulatory interactions between them in the responding epiblast cells, over a period of 12h before definitive neural plate markers like *Sox1* start to be expressed(20). The fine temporal resolution of the GRN (1-2 h) makes it a powerful resource to explore signaling inputs from the node that regulate early steps in the neural induction cascade.

Here we explore the signaling properties of the node. We first explore the expression of signaling molecules by node cells, and reveal that there is considerable cell diversity, with signals associating with particular transcription factor signatures, suggesting different subpopulations of signaling cells. We then use the GRN to explore the targets of each of 22 signals when administered alone. We identify several GRN components that are regulated by signals not previously explored, but not by any of the major known signaling inputs of neural induction (FGF, BMP-inhibition, Wnt-inhibition, Shh and retinoids). Overall, our study suggests that the organizer is a collection of different cell subpopulations with distinct signaling properties, that cooperate to assemble the combination of signals that achieve the inducing and patterning properties of the node.

## Results

### The organizer expresses many signaling molecules in a heterogeneous way

The tip of the chick primitive streak has full neural inducing activity from stages(21) (HH) 3^+^-4, after which the node starts to lose its inducing ability(22-25). We began this study by surveying the signaling molecules expressed at the tip of the streak/Hensen’s node in the chick embryo using transcriptomics of whole nodes, separated node regions and single cells at stage HH3^+^/4 (Fig. 1 A, B). Re-analysis of previous microarray datasets(26) revealed 168 putative secreted signaling molecules expressed at the tip of the primitive streak more strongly than at its opposite extreme (Supplementary Table S2). To refine this, we performed RNA-sequencing (RNAseq) of nodes, subdividing it into medial/anterior, posterior-left and posterior-right sides at HH3^+^/4^-^ (Figure 1B). Supplementary Table S3 shows details of 99 genes encoding putatively secreted proteins that are expressed at a level >10 fragments per million kilobases (fpkm) in at least one of those three samples. Whole mount in situ hybridization for 18 of these transcripts confirms these results (Supplementary Fig. S1) as well as revealing other features. For example some genes, including THPO, are expressed largely in the dorsal parts of the node, in a ‘salt-and-pepper’ pattern within node sub-regions, with small groups of neighboring cells having high or low levels of expression (Fig. 1F, Supplementary Fig. S1R) (see also references(27, 28)).

**Figure 1.**
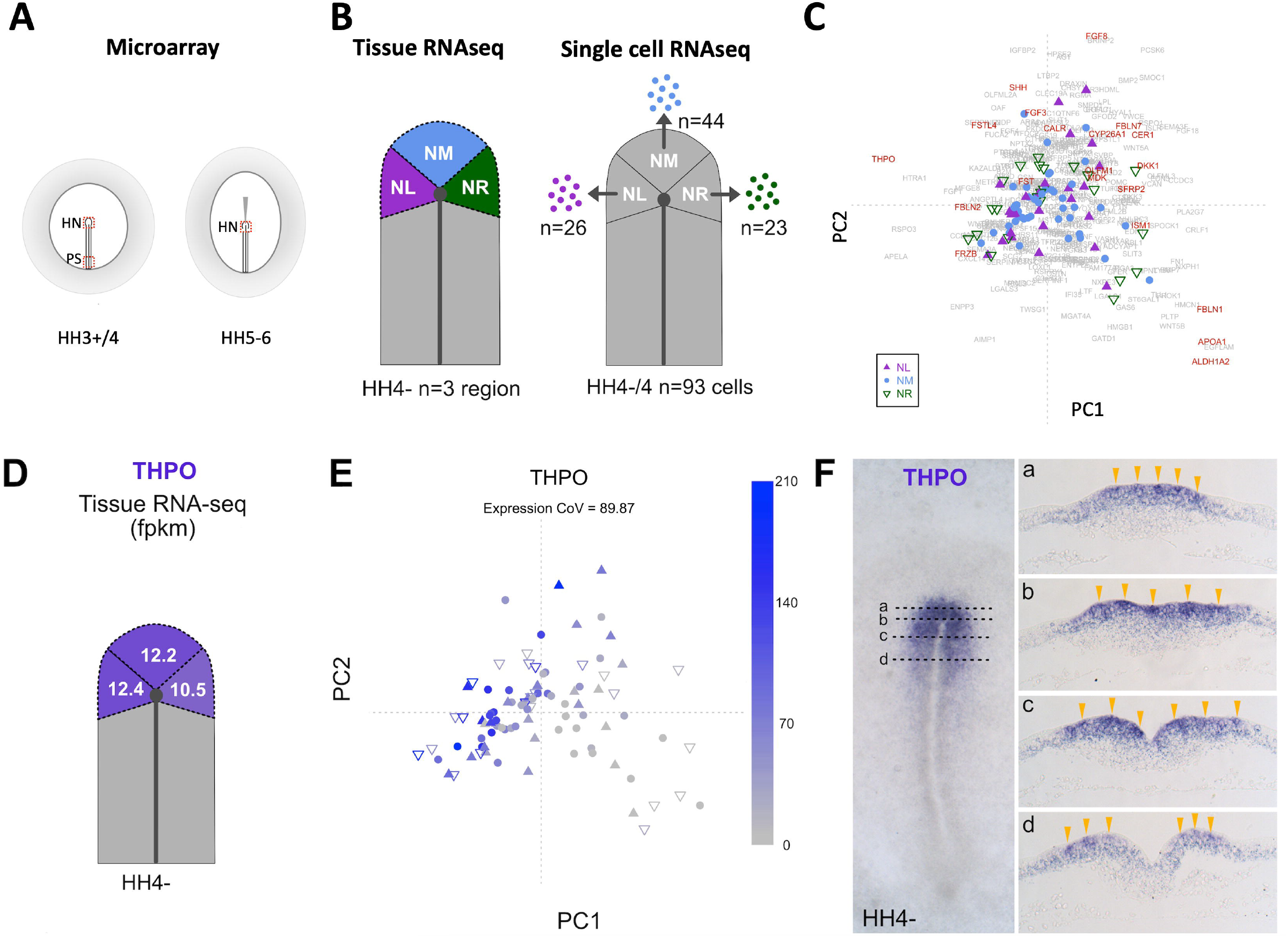
Heterogeneous expression of signaling molecules in the organizer. **A**. Regions sampled for RNA analysis by microarrays, including HN (Hensen’s node), and PS (Posterior primitive streak) at stages HH3+/4 and HH5-6. **B** (left). Tissue RNAseq was performed on three sub-regions (NL: node left; NM: anterior/median node; NR: node right) of the node at HH4-. **B** (right). Single-cell RNAseq (SMART-Seq v4) was performed on 93 cells from the same three sub-regions at HH4-/4. **C**. Cells in the node clustered by their expression of secreted molecules at >10 fpkm in ≥5 cells. The secreted molecules in red font are discussed in Results section. **D**.-**F**. Expression of THPO by tissue RNAseq (D), single cell RNAseq (E) and in-situ hybridization (F); sections on the right at the levels shown in the whole mount on the left.

To obtain a clearer idea of the degree of heterogeneity of expression of signaling molecules, we performed RNAseq from 93 single cells, individually harvested from the medial/anterior and left- and right-posterior sides of HH4^-^/4 nodes (Fig. 1B). Coefficient of variation and principal component analysis revealed that while some secreted molecules are expressed fairly uniformly in these cells (e.g. MDK, MIF, PPIA), others show considerable variation, even within the same region of the node at the same stage (e.g. APOA1, CER1, DKK1, FGF3, and THPO; Fig. 1C) (Supplementary Fig. S2, Supplementary Table S4), revealing patterns not evident from tissue-level RNAseq (Fig. 1 D, E).

### Expression of signaling molecules correlates with transcription factor signatures

To understand whether different cells in the node that express different subsets of signaling molecules have distinct cell identities (different transcription factor expression signatures), we calculated the Pearson’s correlation coefficient between each secreted molecule and transcription factors expressed in the same node cells (fpkm>10 in ≥5 cells), selecting the pairs that have correlation | r | > 0.4 (Fig. 2). Different subsets of secreted molecules were found to be co-expressed with specific combinations of transcription factors. There appear to be at least three broad classes of cells. One subset of cells expresses signals including RSPO3, THPO, APELA and HTRA1, together with a characteristic set of transcription factors including TCF7L1, DNMT3A and 3B, and MTA, but negatively correlates with expression of GATA6, GATA5, SOX17, TCF7L2 and SP5 (Fig. 2 A, C-F). A second, relatively small subset of cells that expresses the signals WNT8A and RGMA, is associated with high expression of ZIC2, RORA, LEF1, KLF5 and PRDM1 (Fig. 2 A, G, L). A third subclass of cells expresses signals such as SFRP1, SLIT1, FGF8, FGF18, RSPO1, FSTL1 and CER1 along with transcription factors including SOX17/GATA5/GATA6, suggesting that they are endoderm precursors (previously suggested to be a critically important source of neural inducing signals(29)) (Fig. 2 A, H-K, M-U). Importantly, the heterogeneity is not due to contamination by non-node cells, because all cells analyzed expressed the node markers GSC and FOXA2 (Supplementary Table S5).

**Figure 2.**
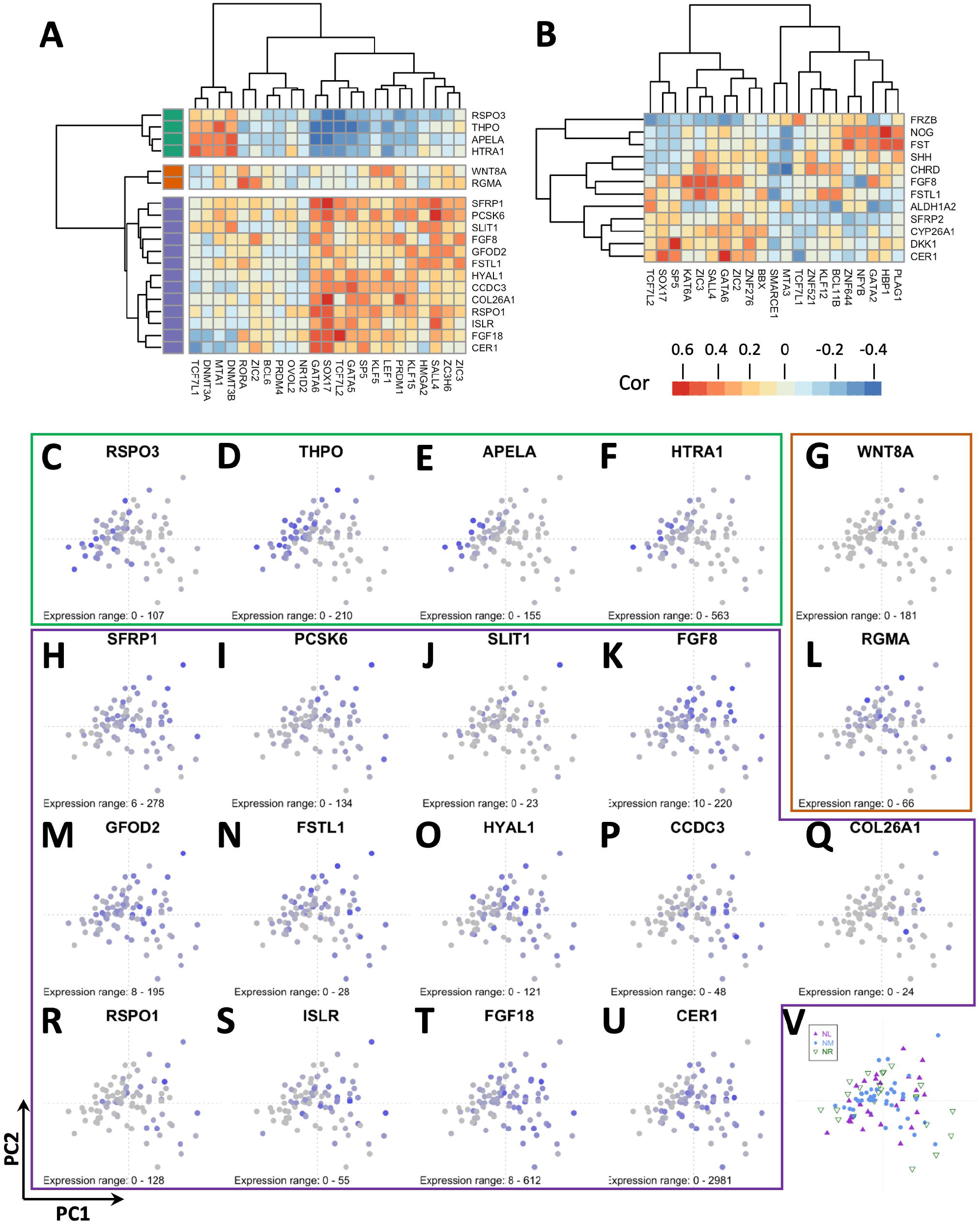
Heatmap showing correlation between secreted molecules and transcription factors. **A**. Correlation between secreted molecules (y axis) and transcription factors (x axis) with Pearson’s correlation coefficient (cor) | r | > 0.4. **B**. Secreted molecules that are known to be involved in neural induction were selected to calculate cor with transcription factors. Transcription factors with | r | > 0.25 relative to any of the known secreted molecules are shown in the heatmap. **C**.-**V**. Principal component analysis based on the expression level of secreted molecules and transcription factors in A. The expression level of RSPO3 (C), THPO (D), APELA (E), HTRA1 (F), WNT8A (G), SFRP1 (H), PCSK6 (I), SLIT1 (J), FGF8 (K), RGMA (L), GFOD2 (M), FSTL1 (N), HYAL1 (O), CCDC3 (P), COL26A1 (Q), RSPO1 (R), ISLR (S), FGF18 (T), and CER1 (U) in each cell (a data point in PCA) are colour-coded from blue (higher expression level) to grey (low or no expression). The regions (NL: node left; NM: node middle; NR: node right as shown in Fig. 1B) from which cells were collected are shown in V.

We also analyzed the expression of some well-characterized signals previously implicated in neural induction (such as CHRD, NOG, FSTL1, FGF8, Wnt-inhibitors and SHH) using a Pearson’s correlation | r | > 0.25. These are also expressed in different subsets of cells (Fig. 2B). For example, the Wnt-antagonists FRZB, DKK1 and SFRP2 are expressed in different subpopulations of cells even though they all play a role as WNT antagonists, and while the BMP inhibitor CHRD is expressed in many cells, two other very well-studied BMP antagonists, NOG and FST, are only expressed strongly in a very small set of cells that do not express CHRD, characterized by expression of transcription factors like PLAG1, HBP1, GATA2, NFYB and ZNF644 (Figure 2B; Supplementary Fig. S3).

Together, these findings indicate that at stages HH3^+^/4, the node is subdivided into domains and that, even within each region, there is heterogeneity in the expression of signaling molecules and transcription factors, suggesting that the node is a collection of distinct sub-populations of cells with different molecular, and presumably different functional, properties.

### Functional analysis of secreted factors expressed in the node

For an initial indication of whether some of the signaling molecules expressed in the node could play a role in neural induction of competent epiblast, we obtained ligand-receptor pairs from CellChatDB (www.cellchat.org)(30), mapping the human homologs of signaling molecules expressed in any region of the chick node at stage 3^+^/4^-^ and receptors expressed in competent area opaca epiblast. The results are shown in Supplementary Table S6; many signaling-receptor pairs are expressed appropriately at significantly high levels in the two tissues. The most prominent ligands include FGF8 and FGF18, MDK, Ephrin B2, Semaphorins 4D and 5D and GDF3 (VG1).

A newly described, comprehensive gene regulatory network (GRN) marking the responses of epiblast cells to a graft of the node in time-course with fine resolution(20) offers a tool to explore which parts of the network respond to different signals. Because of the complexity of the neural induction network, we focused on transcriptional responses occurring within the first 5h of exposure to secreted factors. We started by exploring five pathways that had previously been implicated in neural induction either in chick or in other vertebrates: BMP-inhibition, FGF, Wnt-inhibition, Sonic hedgehog and retinoic acid. A bead loaded with the pure factor was grafted to the inner area opaca at HH3^+^/4^-^ (Supplementary Fig. S4B, Supplementary Table S7; see Materials and Methods for details); after 5h incubation, the bead was removed and the adjacent epiblast (∼200μm diameter) collected. Each sample comprised 7-11 pooled explants, and the experiment was performed in triplicate. Similar explants were collected from embryos grafted with vehicle-loaded control beads. Samples were also collected from embryos grafted with a Hensen’s node in the same way, for comparison (Supplementary Fig. S4A). Samples were analyzed by NanoString nCounter analysis for all GRN components (175 genes) and other markers (see Materials and Methods for details of the probes) (381 genes in total). To evaluate the results, we asked whether the factor alone was “sufficient” to change the expression of genes in the same direction (up- or down-regulation) as does a graft of the node, relative to the contralateral epiblast (5h incubation). The results are summarized in Figure 3 A, D, G and Supplementary Table S9, for each of the five pathways. Out of the 142 GRN components that are up- or down-regulated within 5h(20), 74 are appropriately upregulated (induced), and 39 appropriately downregulated (repressed) by at least one of these five signaling pathways within 5h.

**Figure 3.**
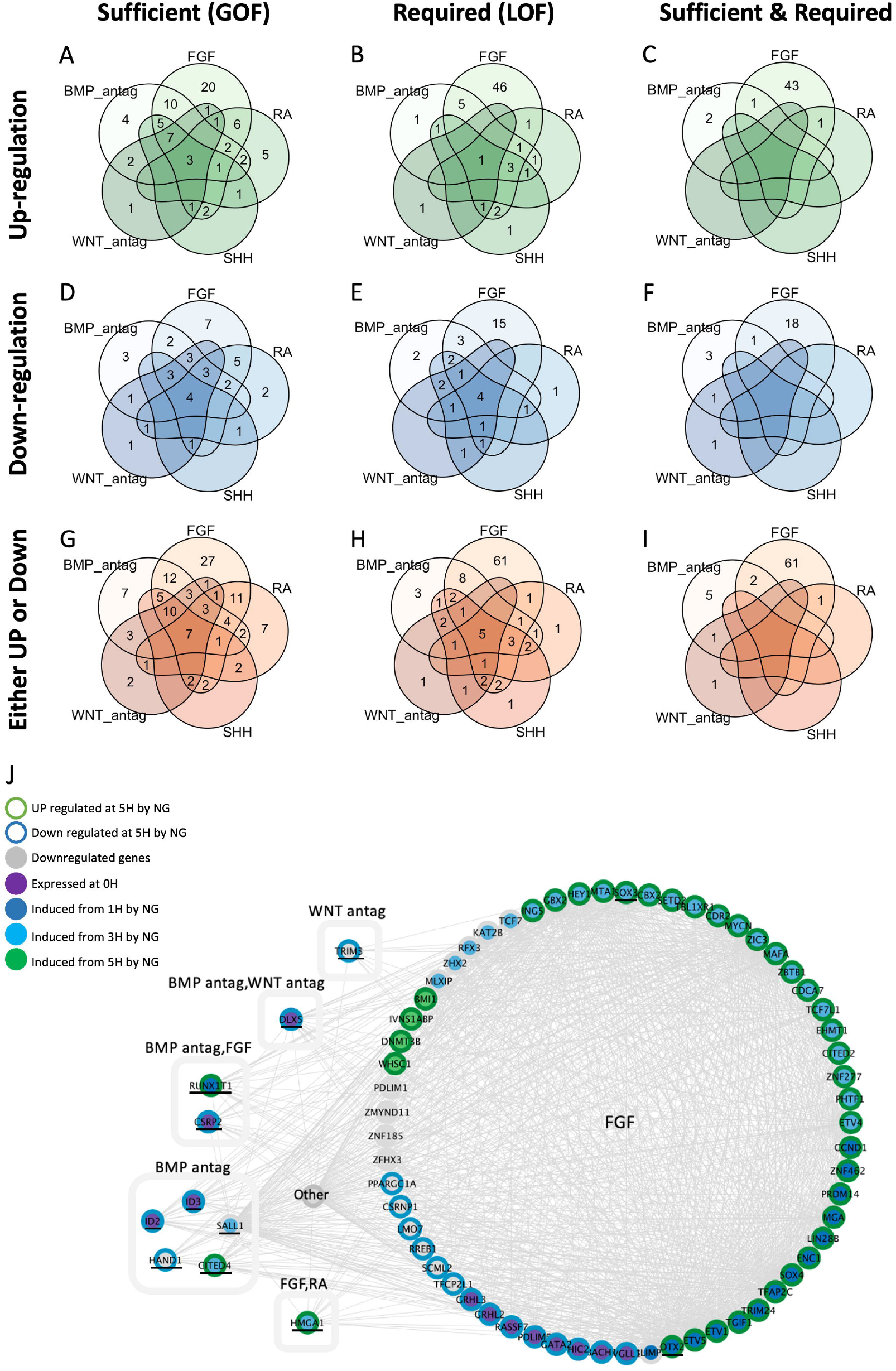
Genes regulated by the five main signaling pathways previously described in the literature. **A.-I**. Venn diagrams showing the numbers of genes regulated by the five signaling pathways previously reported to play a role in neural induction from gain-of-function (“Sufficient”), loss-of-function (“Required”), or both (“Sufficient and Required”). **J**. Network showing the components that are regulated (both “Sufficient and Required” either UP-or DOWN-regulation) by one or more of these main signaling pathways. NG: node graft.

To explore whether these five pathways are required for epiblast cells to respond to a node graft, we performed “loss-of-function” experiments by grafting a node (HH3^+^/4^-^) into a similar region of the area opaca, surrounded by 2-3 beads of an inhibitor of the pathway being explored, or in some cases with the whole host embryo treated with a soluble inhibitor (Supplementary Fig. S4C; Supplementary Table S8; see Materials and Methods for details). After 5h incubation, we removed the node and beads and collected explants of adjacent epiblast to assess changes in gene expression as described above, compared to similar explants from embryos grafted with a node alone, or with the node along with epiblast/node exposed to vehicle (Supplementary Fig. S4E). We scored samples showing at least 20% reduction, or 20% increase in the expression, respectively, of genes that are induced or repressed by a node graft alone. These results are summarized in Fig. 3 and Supplementary Table S9.

Of the five pathways, FGF generates the most responses at 5h. Sixty-one GRN components are regulated by FGF alone and their induction or repression by a node is suppressed by the loss-of-function treatment (i.e. the pathway is both “necessary” and “sufficient” for regulation of the GRN component in the same direction as in the network) (Fig. 3J). This set of genes includes pre-neural plate markers like *OTX2* and *SOX3*. Five transcription factors (*ID2, ID3, SALL1, HAND1* and *CITED4*) are regulated by BMP-inhibition alone (again as “necessary and sufficient”), and one by Wnt-antagonism (*TRIM3*). For a few genes, more than one pathway appeared “necessary and sufficient”: two (*CSRP2* and *RUNX1T1*) were regulated by FGF and also by BMP inhibition, one (*DLX5*) could be regulated by BMP- and by Wnt-antagonism, and one (*HMGA1*) by FGF and retinoic acid (Fig. 3B, Supplementary Table S9). Of the 142 GRN components that are differentially expressed during the first 5 h in the network, 71 (50%) are regulated by at least one of these 5 pathways (satisfying “necessary and sufficient” criteria). In other cases, exposure to one of the factors was “sufficient” to mimic a response to a node graft (e.g. Sonic hedgehog upregulated 29 genes, but inhibition of that pathway did not abolish the response of some of these genes to a node graft (e.g. *SETD2*; Fig. 3A)). However, a significant number (13) of GRN components is not regulated (satisfying either “necessary”, or “sufficient”, or both conditions) by any of these five pathways during the first 5 h of exposure to the signal. Moreover, 29/142 of the 5h GRN components were not regulated in gain-of-function experiments (“sufficient”) for any of the five pathways.

The initial microarray analysis comparing Hensen’s node to the posterior primitive streak identified 169 putative secreted molecules (Fig. 1A, Supplementary Table S2) enriched in the node at appropriate stages of development to be involved in neural induction(26). Subsequent RNAseq analysis identified 177 transcripts encoding putative secreted molecules with a minimum abundance of at least 10 fpkm in tissue preparations of node RNA, or 10 fpkm by scRNAseq in at least 30 cells (Supplementary Table S3, S4; Supplementary Fig. S6). Could some of these signals regulate GRN components not affected by any of the five major pathways? To select suitable candidates, we triaged this set of factors: first, we used information about their expression (for 80 genes whose expression had either been published, or was analyzed by in situ hybridization in our lab), eliminating factors that were either ubiquitous or expressed within the area opaca itself. Second, we eliminated integral membrane proteins and structural components of the extracellular matrix. Finally, we selected the remaining genes whose protein products are available commercially. This generated a list of 27 proteins (10 related to the 5 pathways previously implicated in neural induction as discussed above, and 17 novel secreted molecules) (Supplementary Table S1). To test their ability to mimic a node graft in terms of regulating the expression of components of the GRN, we used AffiGelBlue or Heparin acrylic beads (Supplementary Table S7) grafted alone in the competent region of the area opaca as described above, and compared the responses of equivalent cells that had been exposed to a node graft. NanoString analysis was used to assess the responses after 5h exposure. The full list of factors is shown in Supplementary Table S1. Briefly, we find that 30 out of 381 genes on the NanoString probe set respond to at least one of the new proteins (up- or down-regulation in the same direction as triggered by the node graft), including 16 GRN components that are not regulated by any of the five major pathways previously implicated in neural induction (for example *CEBPB, HOXA1, IRX1* and *SOX11*; Fig. 4; Supplementary Fig. S5). Notable among these signals are some proteins that function as neurotransmitters or have other roles at later stages of neural development, such as members of the SLIT, Olfactomedin and Somatostatin families, as well as Fibulins, Thrombopoietin and enzymes involved in the production of some prostaglandins.

**Figure 4.**
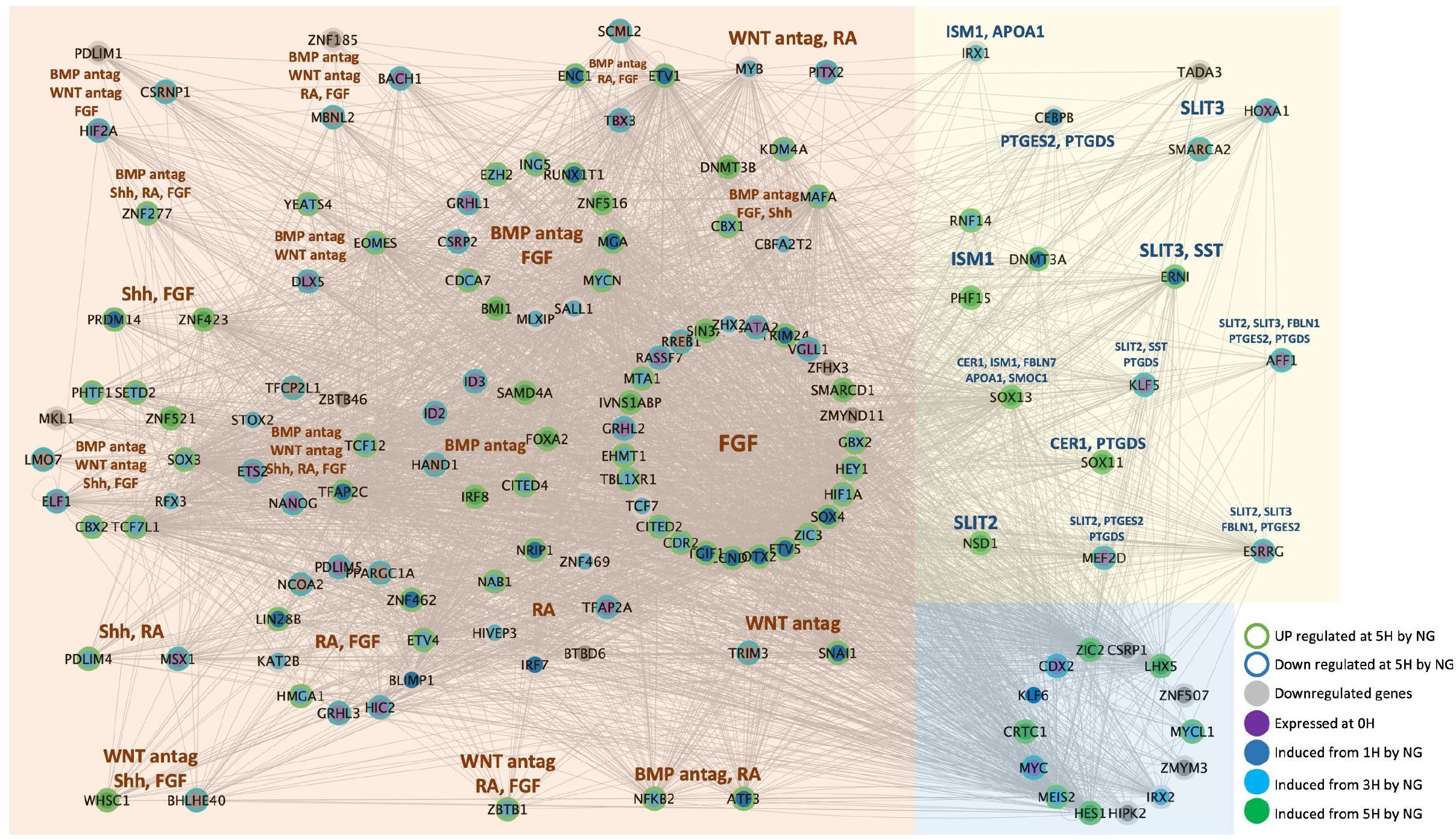
Neural Induction GRN components regulated (“sufficient”) by signaling factors in the first 5 hours. Network components shown in the circle on the left (orange box) are regulated by at least one of the five main signaling pathways previously described; components on the right (yellow box) are regulated by novel secreted molecule(s) studied here. Components in the blue box were not regulated by any of the signaling inputs studied. The color of the components represents the initial time point of expression in the GRN(20); the color of the edge shows whether the component is regulated by a node graft at 5h by a node graft.

Together, our results suggest that many (113/142) GRN components whose expression changes in the first 5h following a graft of the organizer (Hensen’s node) can be accounted for by the activity of FGF, BMP-inhibition, Wnt-inhibition and to a lesser extent RA and Shh, and that 11 of the 17 novel secreted factors expressed in the node at the appropriate stage that have been tested can regulate 16 of the remaining 29 genes (Supplementary Table S11).

## Discussion

Previous experiments in the chick involving an organizer graft and its removal after increasing periods of time established that 12-13h contact with the organizer are required for non-neural ectoderm cells to be fully directed to a neural plate fate(15, 24, 25, 31, 32). A variation of this experiment, in which the organizer was removed and replaced by Chordin-secreting cells, demonstrated that 5h signaling by the node are required for the ectoderm to become responsive to inhibition of BMP(15), suggesting that the process is initiated by signals other than BMP inhibition. FGF turned out to be a critical signal for this(12-14, 33-36). The first 5h of signals from the organizer are therefore crucial for initiating the process of neural induction(20, 36, 37). Consistent with these findings, here we find that FGF accounts for many (61/142) genes whose expression changes in response to the node within the first 5h (satisfying both “necessary” and “sufficient” criteria), whereas BMP inhibition alone only regulates 4/142 genes (Fig. 3).

We also previously reported that even when epiblast is first exposed to FGF for 5h, followed by strong BMP inhibition, the combination is still not sufficient to induce a stable neural plate(13, 14, 36), suggesting that other pathways are involved. Analysis of the responses to three other signals previously implicated in neural induction (Wnt-inhibition, Sonic hedgehog and retinoic acid) still leaves 29/142 (20%) of the GRN genes regulated by the node in the first 5h unaccounted for. Here we show that 16 of these genes are regulated by 11 signals that had not previously been described as playing a role in this process, and another 6 novel signaling molecules expressed in the node regulate genes that are also regulated by at least one of the 5 pathways previously implicated in neural induction (Supplementary Table S10).

### Heterogeneity in the node: what is “the organizer”?

Our study reveals considerable heterogeneity within the organizer. Several signaling molecules are expressed in cells with characteristic transcription factor expression signatures (some correlated positively, some negatively), strongly suggesting the existence of distinct sub-populations of signaling cells. In turn, this raises the possibility that the organizer could be more accurately described as a ***cell collective***, whose members cooperate by providing complementary signals that together result in the induction and patterning of the CNS. Some of these cell sub-populations are localized to particular regions of the node (e.g. left or right, anterior-median or posterior-lateral), but others appear to be randomly interspersed with other cell populations in a salt-and-pepper fashion. A previous study comparing the neural inducing activities of different sub-regions of the node (epiblast versus deep, anteromedian versus posterolateral) reported that neural induction is mainly associated with the epiblast portion of all domains, and the deep portion of the antero-median sector, whereas deep posterolateral cells have less inducing ability(38). Interestingly, the posterolateral domain of the regressing node at HH8 is the region that contains the niche(s) for resident, self-renewing cells that generate progeny associated with caudal elongation(27, 39), whereas all regions of the node contain more transient populations of cells. A view is therefore starting to emerge of the node/organizer as a location rather than a fixed population of cells, within which there are highly dynamic changes in cell composition(14, 27, 33, 39-42). We now show that this includes changes in signaling properties, which define the inducing and patterning functions of this region by the collective, cooperative action of these sub-populations.

Although the original idea proposed by Otto Mangold that there could be separate organizers responsible for inducing different regions of the neural plate (head, trunk and tail organizers, perhaps many more)(43-45) now seems very unattractive(46), it is conceivable that the node does contain separate populations of cells with specific regional inducing functions that change over time, which could account for the induction of a fully patterned nervous system. One observation that supports this speculation is that an “older” (stage HH4^+^/5) Hensen’s node transplant into the area opaca, which normally can no longer induce anterior neural plate in this assay, is recombined with the cells that have left the node between HH4 and 4^+^ (the prechordal mesendoderm), the combination can now induce a full neural plate including the forebrain(47, 48). One prediction made by this proposal is that at least some of these populations should change between stage HH4 (when the node can induce a complete CNS) and HH5-6 (when it can only induce more caudal CNS(22, 24, 29, 49, 50)).

### Conclusions

In the process of neural induction by a grafted node, it is likely that many signals act both simultaneously as well as sequentially and that the process continues beyond 5h, as suggested by node removal experiments (indicating a requirement for signals for 12-13h)(15). Our transcriptome analysis of the node suggests that as development proceeds, the precise combination of secreted proteins it produces changes. It is therefore impossible to design experiments to assess the contributions of all combinations of these signals, even if given together, in a manner that resembles the complex signaling activity of the organizer.

It should also be mentioned that we have also opted to use only a single concentration of each signal to test. To arrive at this concentration for each factor, we were guided by published information, product data sheets and when possible, an independent “positive control” experiment testing various concentrations before choosing an efficient but realistic concentration to use on the embryo (see Materials and Methods and Supplementary Table S7, S8 for details).

Despite these caveats, which suggest even more complexity, our study reveals that rather than a static population of unique cells (as suggested from the uniform expression of organizer markers like GSC and FOXA2) that emit one or a few signals responsible for its inducing functions(51), the organizer comprises a mixture of cell sub-populations that can be defined by transcriptional signatures, which emit different signals that cooperatively regulate a complex network of genetic responses that initiate induction of the Central Nervous System.

## Materials and Methods

The following is a summary of the methods used. Full details are given in Supplementary Information.

### Embryos and mRNAseq

Egg incubation, staging, Hensen’s node transplants, tissue collection for tissue RNAseq and in situ hybridization were performed as previously described(21, 24, 26, 27). Tissue RNAseq was done on 3 subregions from HH4-nodes (NL-node left; NM-node middle; NR-node right) as illustrated in Fig. 1B. The tissue RNAseq raw data have been deposited in European Bioinformatics Institute (EBI) Array Express (E-MTAB-14510). scRNAseq was performed on 93 cells from HH4-/4 nodes. Individual cells were collected from the same subregions of the node (Fig. 1B) and scRNAseq performed as previously described(27). The scRNAseq raw data have been deposited in EBI Array Express (E-MTAB-14518).

To select secreted molecules that may be associated with neural induction, we applied the following criteria to the RNAseq data: 1) genes annotated as “secreted proteins” in the Human Protein Atlas (www.proteinatlas.org), 2) the presence of a signal sequence identified by SignalP 4.1, 3) FPKM expression level >10 in at least one of the node tissue samples (tissue RNAseq) or FPKM >10 in at least 30 collected cells (scRNAseq). The resulting secreted molecules are listed in Supplementary Tables S3 and S4.

### Bead implantation and chemical treatments

The factors or chemical agents used, their commercial source and concentration used are detailed in Supplementary Table S7. The type of bead or method of delivery used is also shown in the table. As a rule of thumb, we used AG1×2-formate-modified beads for agents that are water-insoluble but soluble in DMSO or other organic solvents (such as retinoic acid), Heparin-acrylic beads for FGF and other heparin-binding proteins, and AffiGel-Blue for other water-soluble proteins. Beads were loaded overnight at 4°C for at least 12 h and then rinsed twice briefly with PBS to remove excess factor before implantation into the inner area opaca at stage 3+/4-. For some chemical agents dissolved in DMSO, whole embryos were soaked in a dilute solution for 40 min then rinsed briefly prior to culture with the chemical diluted in the albumin underlying the embryo. Loss-of-function assays were conducted in a similar way but together with a graft of HH3+/4-node; details are given in Supplementary Table S8.

### NanoString tissue collection and processing

Epiblast that had been exposed to a node graft and/or beads was dissected as previously described (20, 37). Seven (up to eleven) explants (about 200μm in diameter) were pooled for each sample, and all experiments conducted as biological triplicates (three independent pools per condition). NanoString experiments were run on the nCounter Analysis System following NanoString guidelines, using a custom codeset of 386 probes as described previously(20, 37).

### NanoString expression data analysis

Each experimental condition (either GOF, LOF or node graft) was compared to its own control (Supplementary Tables S6 and S7) and calculated for mean, fold change (FC) and p value (two-sided t-test). To determine whether a factor alone is sufficient to regulate the expression of a GRN component in the same way as a node graft (either up- or down-regulation), we checked changes of the gene expression from GOF and node graft experiments using the following criteria (Supplementary Fig. S4D): 1) comparing GOF of a secreted molecule with its control p value < 0.05; 2) GOF and node graft experiments compared with their own controls, with thresholds: FC >= 1.2 for upregulated genes, or FC <= 0.8 for down-regulated genes; 3) gene expression mean v5alues >40 fpkm in both GOF and node graft induced tissue samples for upregulated genes, and >40 fpkm in GOF-control and node-graft-lateral-control (uninduced contralateral) samples for down-regulated genes.

To determine whether a secreted factor is required for a GRN component to be differentially expressed, we checked for changes in gene expression from LOF and node graft experiments with criteria (Supplementary Fig. S4E): 1) comparing LOF of a secreted molecule (inhibitor of the secreted molecule plus a grafted node) with its own control p value < 0.05; 2) genes that require the secreted factor for upregulation, LOF test compared with its control FC <= 0.8 and node graft compared with its lateral control FC >= 1.2; for genes requiring the secreted factor for downregulation, LOF test compared with its control FC >= 1.2 and node graft compared with its control FC <= 0.8; 3) upregulated genes with mean expression values >40 fpkm, downregulated genes with mean expression values >40 in lateral control sample (uninduced contralateral); 4) expression FC from LOF test at least 20% lower than the FC from the node graft experiment for genes whose upregulation requires the factor, and expression FC from LOF test at least 20% higher than the node graft experiments for genes whose downregulation requires the factor.

### Identification of cell subpopulations from single cell RNAseq data

Secreted molecules and transcription factors were selected based on the gene annotations “secreted proteins” and “transcription factors” in the Human Protein Atlas (www.proteinatlas.org) that are also expressed at fpkm > 10 in at least 5 cells in the scRNAseq data. Pearson’s correlation coefficient (cor) was then calculated for each pair of a secreted molecule and a transcription factor. Pairs with absolute value of cor | r | > 0.4 were selected to generate the heatmap (Fig. 2A) and principal component analysis (PCA) (Fig. 2V). Previously well-characterized secreted molecules involved in neural induction were separately selected to calculate cor values with transcription factors. The pairs were filtered with absolute cor | r | > 0.25 to generate the heatmap (Fig. 2B) and PCA plot (Supplementary Fig. S3A).

## Supporting information

Supplementary information: Detailed Methods

## Acknowledgments

This study was funded by grants from NIH (R01 MH60156) and BBSRC (BB/R003432/1 and BB/K007742/1) to CDS. CDS was a Wellcome Trust investigator (107055/Z/15/Z). KT was a BBSRC quota PhD student. TS, LF and CC were students in the Wellcome Trust funded PhD programme “Developmental and Stem Cell Biology” at UCL.

## Notes

### Competing Interest Statement

The authors have declared no competing interest.

