## Supplementary information: Detailed Methods for "The organizer as a cooperative of signaling cells for neural induction"

**MATERIALS AND METHODS**

*Embryos*

Fertile hen’s eggs (Brown Bovan Gold; Henry Stewart & Co., UK) were incubated at 38°C in humidified incubators and staged according to Hamburger and Hamilton^1^ (HH). All RNAseq analyses of the node were performed on tissue/cells from transgenic cytoplasmic-GFP embryos from The Roslin Institute, Edinburgh. Embryos were harvested and cultured ex ovo using the New technique with modifications as previously described^2,3^. Transplantation of the node was performed as previously described^4-6^.

*RNAseq tissue collection and processing*

Tissue RNAseq was done on 3 subregions from HH4- nodes (NL-node left; NM-node middle; NR-node right) as illustrated in Fig. 1B. Twelve to seventeen pieces were collected for each region. Dissection, RNA-extraction, measurement, library synthesis and sequencing were performed as described previously^7^. The average number of reads per sample was ~24 million (range:15,442,739-32,338,154). The tissue RNAseq raw data have been deposited in European Bioinformatics Institute (EBI) Array Express (E-MTAB-14510).

*Single cell RNA sequencing sample preparation*

scRNAseq was performed on 93 cells from HH4-/4 nodes. Cells were collected from three subregions of the node (NL-node left; NM-node middle; NR-node right) as illustrated in Fig. 1B. Individual cells were isolated and collected as previously described^7^. Cells were prepared for sequencing using the SMART-Seq v4 Ultra Low Input RNA Kit (Takara, # 634892). cDNA purification, measurement, sonication, library preparation and sequencing were performed as previously described^7^. The average number of reads per cell was about 8.9 million (range: 6,604,828-11,804,768). The scRNAseq raw data have been deposited in EBI Array Express (E-MTAB-14518).

*Whole-mount in situ hybridization*

In situ hybridization with digoxigenin (DIG)-labelled riboprobes was carried out as previously described^6,8,9^. Plasmids used: CHRD^10^; DKK1^11^; CER1^12^; FBLN2 (ChEST655K12); FBLN7; FGF8^13^; FSTL4 (ChEST43301); IGF2 (ChEST990L17); ISM1^14^; OLFM1^15^; PTGDS (ChEST223a22); PTGES2 (ChEST195i9); SHH^16^; SLIT2^17^; SLIT3^17^; SMOC1 (ChEST66m21); SST^18^; THPO (ChEST671e4).

*Bead implantation and other chemical treatments*

The factors or chemical agents used, their commercial source and concentration used are detailed in Supplementary Table S7. The type of bead or method of delivery used is also shown in the table. As a rule of thumb, we used AG1X2-formate-modified beads for agents that are water-insoluble but soluble in DMSO or other organic solvents (such as retinoic acid), Heparin-acrylic beads for FGF and other heparin-binding proteins, and Affi Gel Blue for other water-soluble proteins. The effective concentration to be used was arrived at by consulting the literature for each factor. Where possible, this was verified using a specific biological assay for each, also detailed in Supplementary Table S7. Where this was not available, beads were loaded with the maximum factor concentration while maintaining low vehicle concentrations of BSA and DMSO. Beads were loaded overnight at 4°C for at least 12 h and then rinsed twice briefly with PBS to remove excess factor before implantation into the inner area opaca at stage 3+/4-. For some chemical agents dissolved in DMSO, whole embryos were soaked in a dilute solution for 40 min then rinsed briefly prior to culture with the chemical diluted in the albumin underlying the embryo.

Loss-of-function assays were conducted in a similar way but together with a graft of HH3+/4- node; details are given in Supplementary Table S8.

*NanoString tissue collection and processing*

Epiblast that had been exposed to a node graft and/or beads was dissected as previously described^6^. Seven (up to eleven) explants (about 200µm in diameter) were pooled for each sample, and all experiments conducted as biological triplicates (three independent pools per condition). Tissues were dissected in ice-cold PBS and collected on ice. Excess solution was removed using a fine needle and promptly processed by adding 1 μl lysis buffer from the RNAqueous-Micro Total RNA Isolation kit (Thermo Fisher). Tubes were immediately snap-frozen on dry ice and stored at ‑80°C. NanoString experiments were run on the nCounter Analysis System using a custom codeset and following NanoString guidelines. The codeset consisted of 386 probes, described previously^6^.

*Tissue and single cell RNAseq data analysis*

Raw data in FASTQ format underwent quality check using FastQC^19^. Cutadapt^20^ was used to remove low-quality (Phred quality score <20) or ‘N’ bases at the 3’ and 5’ ends, and adaptor sequences. Reads were aligned to the galGal6 genome using TopHat2^21^; alignment rates were 87.6% ± 0.14% (bulk RNAseq data) and 94.0% ± 0.3% (scRNAseq data). Transcripts were counted and normalized using Cufflinks^22^ programs *cuffquant* and *cuffnorm*, respectively.

To select secreted molecules that may be associated with neural induction, we applied the following criteria to the RNAseq data: 1) genes annotated as “secreted proteins” in the Human Protein Atlas ([www.proteinatlas.org](http://www.proteinatlas.org))^23^, 2) the presence of a signal sequence identified by SignalP 4.1^24^, 3) FPKM expression level > 10 in at least one of the node tissue samples (tissue RNAseq) or FPKM >10 in at least 30 collected cells (scRNAseq). The resulting secreted molecules are listed in Supplementary Tables S3 and S4.

*Differential expression analysis by microarrays*

The dataset of Hensen’s node at HH3^+^/4 (HH3^+^/4 HN) and HH5-6 (HH5-6 HN), and posterior primitive streak at HH3^+^/4 (PS) were downloaded from ArrayExpression (accession number E-MTAB-4048). The raw data were first normalised using the probe intensity error method (affy library) and linear fitted using limma library in the R environment (R 3.0). Pair-wise comparison of samples was preformed, including HH3^+^/4 HN versus PS, HH3^+^/4 HN versus HH5-6 HN, HH5-6 HN versus PS, to obtain gene probes that are upregulated in HH3^+^/4 HN using threshold log_2_ scale fold change log_2_(FC) ≥ 2 and adjusted p value < 0.05. The secreted molecules were then selected by verifying the presence of a signal sequence identified by SignalP 4.1.

*Correlation between secreted molecules and transcription factors*

Annotations for secreted molecules and transcription factors were obtained from the Human Protein Atlas ([www.proteinatlas.org](http://www.proteinatlas.org))^23^. The normalised expression data in fpkm from single cell RNAseq was used to extract the expression of each secreted molecule and each transcription factor in turn, by calculating Pearson’s correlation coefficient. oThe cells in each heatmap are clustered by the expression of the secreted molecules and correlated transcription factors using the hierarchical clustering method “ward.D2”. The analysis was performed in the R environment (R 4.3.0).

*NanoString expression data analysis*

Nanostring raw data across different experimental conditions (listed in Supplementary Tables S6 and S7) were integrated using NanoString nSolver 4.0 (“New Multi-RLF Experiment” and “Cross RLF and Batch Calibration” options), with background noise subtracted (geomean of negative control probes), and normalised by the expression of housekeeping genes (geomean of ACTB and GAPDH) as described in the nSolver User documentation. The nSolver (“Custom Text Format Export” function) generated normalised data in text format (.tsv).

The R environment (R 4.3.0) was then used for comparative gene expression analysis. Each experimental condition (either GOF, LOF or node graft) was compared to its own control (Supplementary Tables S6 and S7) and calculated for mean, fold change (FC) and p value (two-sided t-test).

To determine whether a factor alone is sufficient to regulate the expression of a GRN component in the same way as a node graft (either up- or down-regulation), we checked changes of the gene expression from GOF and node graft experiments using the following criteria (Supplementary Fig. S8D): 1) comparing GOF of a secreted molecule with its control p value < 0.05; 2) GOF and node graft experiments compared with their own controls, with thresholds: FC >= 1.2 for upregulated genes, or FC <= 0.8 for down-regulated genes; 3) gene expression mean v5alues >40 fpkm in both GOF and node graft induced tissue samples for upregulated genes, and >40 fpkm in GOF-control and node-graft-lateral-control (uninduced contralateral) samples for down-regulated genes.

To determine whether a secreted factor is required for a GRN component to be differentially expressed, we checked for changes in gene expression from LOF and node graft experiments with criteria (Supplementary Fig. S8E): 1) comparing LOF of a secreted molecule (inhibitor of the secreted molecule plus a grafted node) with its own control p value < 0.05; 2) genes that require the secreted factor for upregulation, LOF test compared with its control FC <= 0.8 and node graft compared with its lateral control FC >= 1.2; for genes requiring the secreted factor for downregulation, LOF test compared with its control FC >= 1.2 and node graft compared with its control FC <= 0.8; 3) upregulated genes with mean expression values >40 fpkm, downregulated genes with mean expression values >40 in lateral control sample (uninduced contralateral); 4) expression FC from LOF test at least 20% lower than the FC from the node graft experiment for genes whose upregulation requires the factor, and expression FC from LOF test at least 20% higher than the node graft experiments for genes whose downregulation requires the factor.

*Ligand-receptor analysis*

Human ligand and receptor annotations were obtained from CellChatDB^25^. Ligand and receptor expression values were extracted from tissue RNAseq HH4- node (left, middle and right) and responding tissue (non-neural extraembryonic ectoderm at 0 h from E-MTAB-10409), respectively.

*Network analysis*

Regulatory interactions between transcription factors that are differentially expressed during the first 5h in the GRN for neural induction were obtained from our previous study^6^. A total of 142 GRN components are differentially expressed during the first 5h after a node graft. Transcription factors that are regulated by the same signalling proteins/pathways (sufficient and required by main signalling pathways, or sufficiently regulated by one or more novel signalling molecules) were grouped for display using Cytoscape^26^.

1	Hamburger, V. & Hamilton, H. L. A series of normal stages in the development of the chick embryo. *J Morphol* **88**, 49-92 (1951).

2	New, D. A. T. A New technique for the cultivation of the chick embryo in vitro. *J Embryol Exp Morphol* **3**, 326-331, doi:10.1242/dev.3.4.326 (1955).

3	Stern, C. D. & Ireland, G. W. An integrated experimental study of endoderm formation in avian embryos. *Anat Embryol (Berl)* **163**, 245-263, doi:10.1007/BF00315703 (1981).

4	Storey, K. G., Crossley, J. M., De Robertis, E. M., Norris, W. E. & Stern, C. D. Neural induction and regionalisation in the chick embryo. *Development* **114**, 729-741 (1992).

5	Streit, A. & Stern, C. D. Operations on primitive streak stage avian embryos. *Methods Cell Biol* **87**, 3-17 (2008).

6	Trevers, K. E. *et al.* A gene regulatory network for neural induction. *Elife* **12**, e73189, doi:10.7554/eLife.73189 (2023).

7	Solovieva, T., Lu, H. C., Moverley, A., Plachta, N. & Stern, C. D. The embryonic node behaves as an instructive stem cell niche for axial elongation. *Proc Natl Acad Sci U S A* **119**, doi:10.1073/pnas.2108935119 (2022).

8	Streit, A. & Stern, C. D. Combined whole-mount in situ hybridization and immunohistochemistry in avian embryos. *Methods* **23**, 339-344. (2001).

9	Trevers, K. E. *et al.* Neural induction by the node and placode induction by head mesoderm share an initial state resembling neural plate border and ES cells. *Proc Natl Acad Sci U S A* **115**, 355-360, doi:10.1073/pnas.1719674115 (2018).

10	Streit, A. *et al.* Chordin regulates primitive streak development and the stability of induced neural cells, but is not sufficient for neural induction in the chick embryo. *Development* **125**, 507-519 (1998).

11	Foley, A. C., Skromne, I. & Stern, C. D. Reconciling different models of forebrain induction and patterning: a dual role for the hypoblast. *Development* **127**, 3839-3854. (2000).

12	Zhu, L. *et al.* Cerberus regulates left-right asymmetry of the embryonic head and heart. *Current Biology* **9**, 931-938 (1999).

13	Streit, A., Berliner, A. J., Papanayotou, C., Sirulnik, A. & Stern, C. D. Initiation of neural induction by FGF signalling before gastrulation. *Nature* **406**, 74-78. (2000).

14	Alev, C. *et al.* Transcriptomic landscape of the primitive streak. *Development* **137**, 2863-2874, doi:10.1242/dev.053462 (2010).

15	Barembaum, M., Moreno, T. A., LaBonne, C., Sechrist, J. & Bronner-Fraser, M. Noelin-1 is a secreted glycoprotein involved in generation of the neural crest. *Nat Cell Biol* **2**, 219-225, doi:10.1038/35008643 (2000).

16	Levin, M., Johnson, R. L., Stern, C. D., Kuehn, M. & Tabin, C. A molecular pathway determining left-right asymmetry in chick embryogenesis. *Cell* **82**, 803-814, doi:10.1016/0092-8674(95)90477-8 (1995).

17	Vargesson, N., Luria, V., Messina, I., Erskine, L. & Laufer, E. Expression patterns of Slit and Robo family members during vertebrate limb development. *Mech Dev* **106**, 175-180, doi:10.1016/s0925-4773(01)00430-0 (2001).

18	Lleras-Forero, L. *et al.* Neuropeptides: developmental signals in placode progenitor formation. *Dev Cell* **26**, 195-203, doi:10.1016/j.devcel.2013.07.001 (2013).

19	Wingett, S. W. & Andrews, S. FastQ Screen: A tool for multi-genome mapping and quality control. *F1000Res* **7**, 1338, doi:10.12688/f1000research.15931.2 (2018).

20	Martin, M. Cutadapt removes adapter sequences from high-throughput sequencing reads. *EMBnet.journal* **17**, doi:10.14806/ej.17.1.200 (2011).

21	Kim, D. *et al.* TopHat2: accurate alignment of transcriptomes in the presence of insertions, deletions and gene fusions. *Genome Biol* **14**, R36, doi:10.1186/gb-2013-14-4-r36 (2013).

22	Trapnell, C. *et al.* Differential analysis of gene regulation at transcript resolution with RNA-seq. *Nat Biotechnol* **31**, 46-53, doi:10.1038/nbt.2450 (2013).

23	Uhlen, M. *et al.* The human secretome. *Sci Signal* **12**, eaaz0274, doi:10.1126/scisignal.aaz0274 (2019).

24	Nielsen, H. Predicting secretory proteins with SignalP. *Methods Mol Biol* **1611**, 59-73, doi:10.1007/978-1-4939-7015-5_6 (2017).

25	Jin, S. *et al.* Inference and analysis of cell-cell communication using CellChat. *Nat Commun* **12**, 1088, doi:10.1038/s41467-021-21246-9 (2021).

26	Shannon, P. *et al.* Cytoscape: a software environment for integrated models of biomolecular interaction networks. *Genome Res* **13**, 2498-2504, doi:10.1101/gr.1239303 (2003).
