## Supplementary figures and images for "The organizer as a cooperative of signaling cells for neural induction"

### Supplementary Figure S1

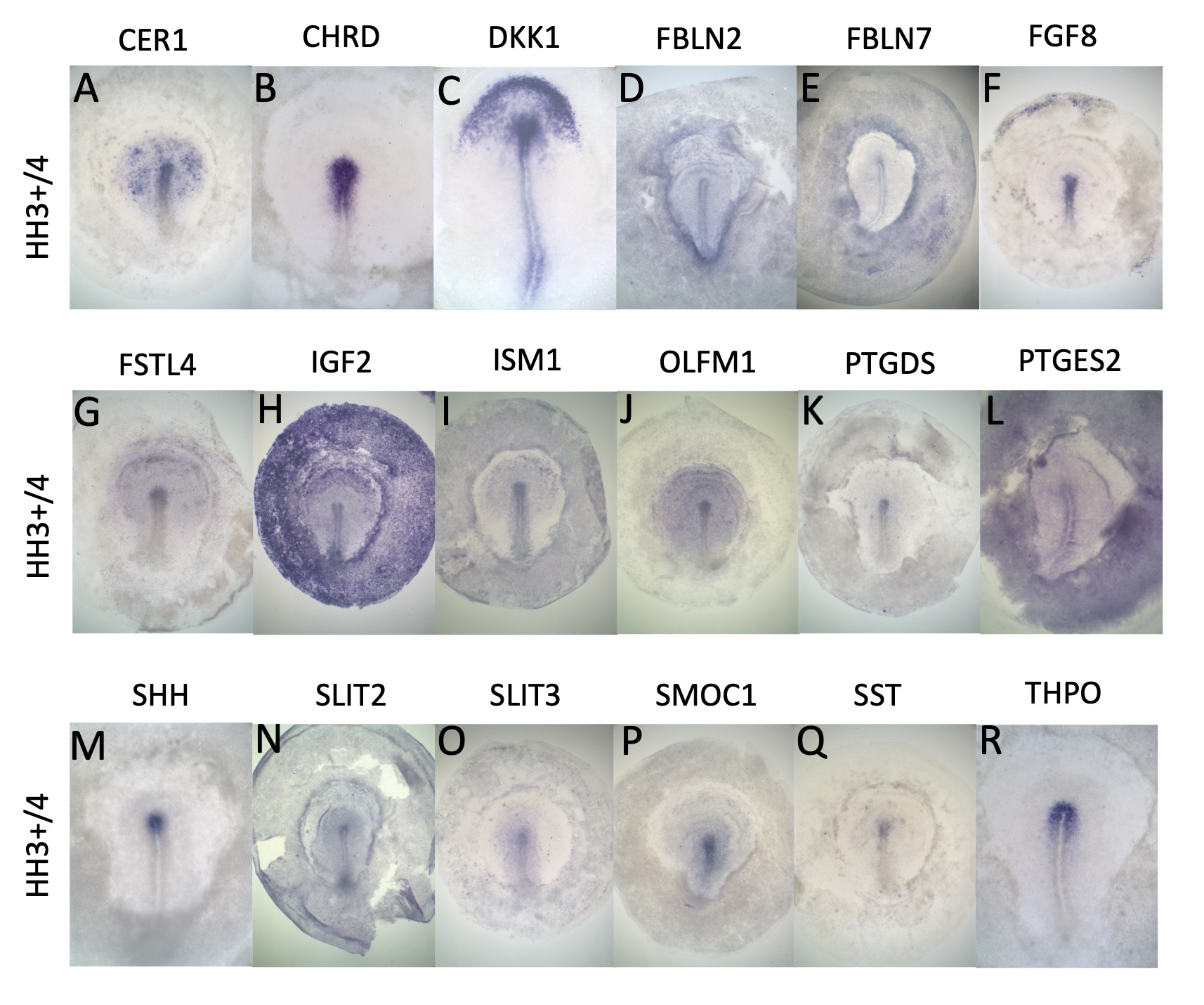

### Supplementary Figure S2

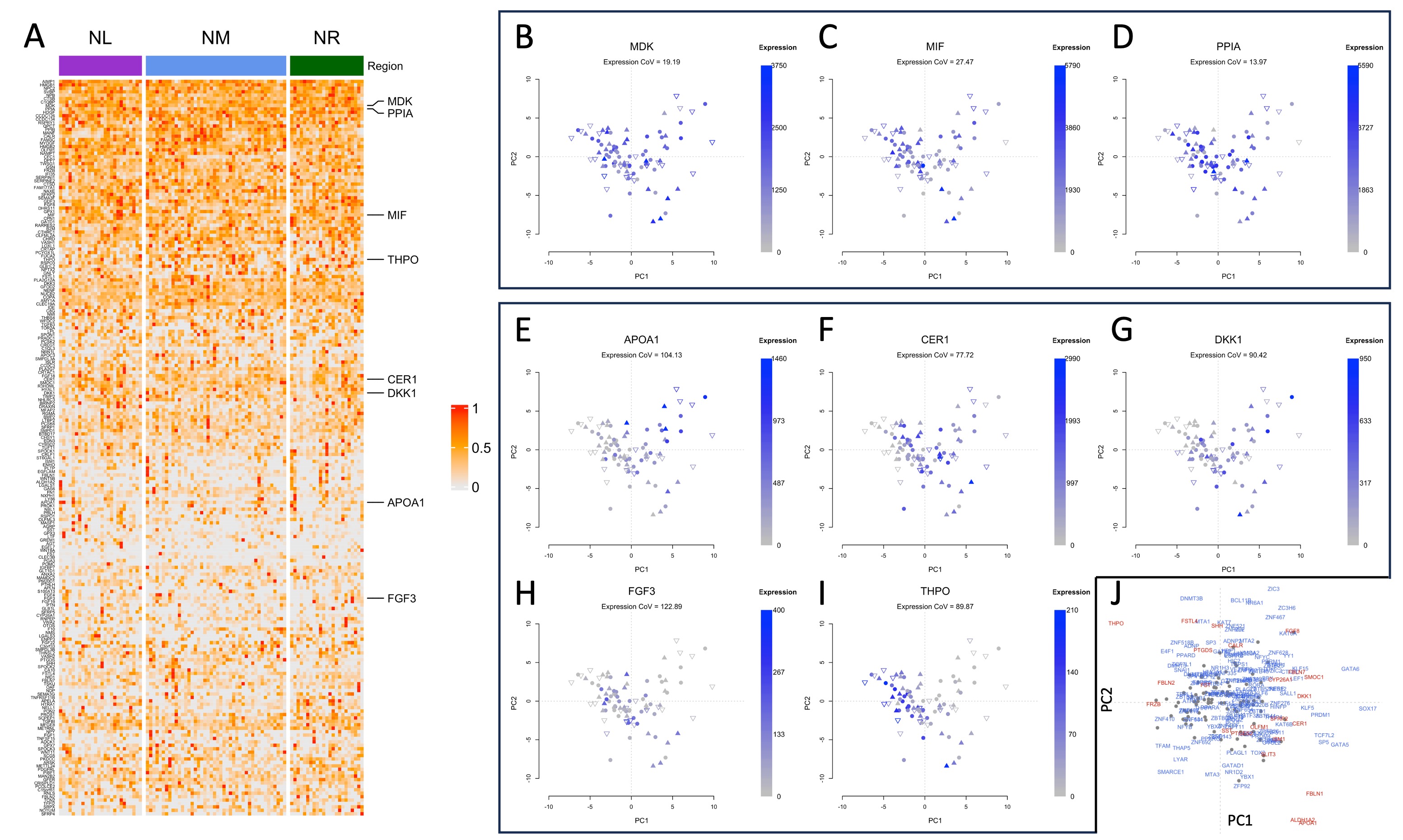

### Supplementary Figure S3

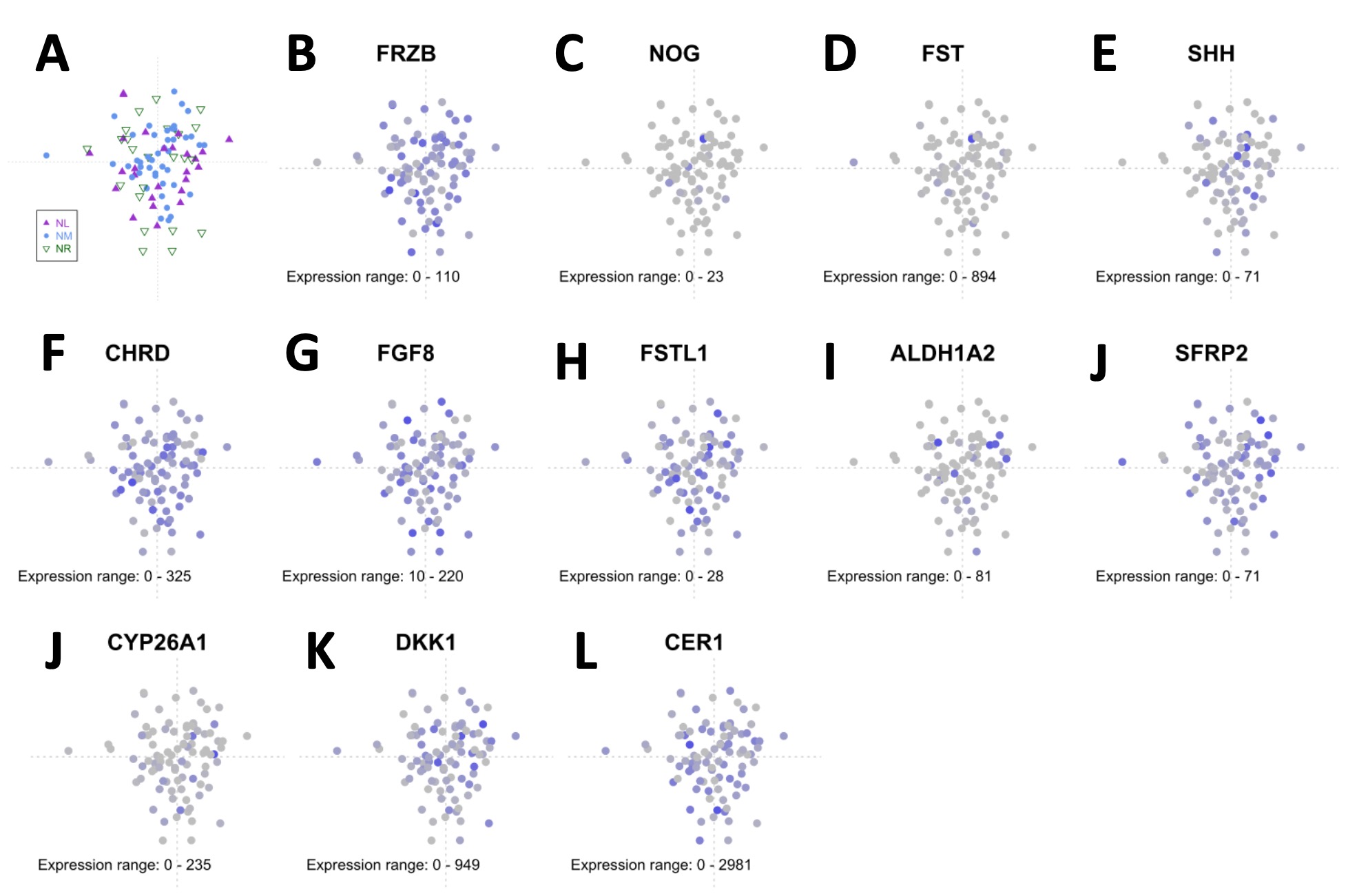

### Supplementary Figure S4

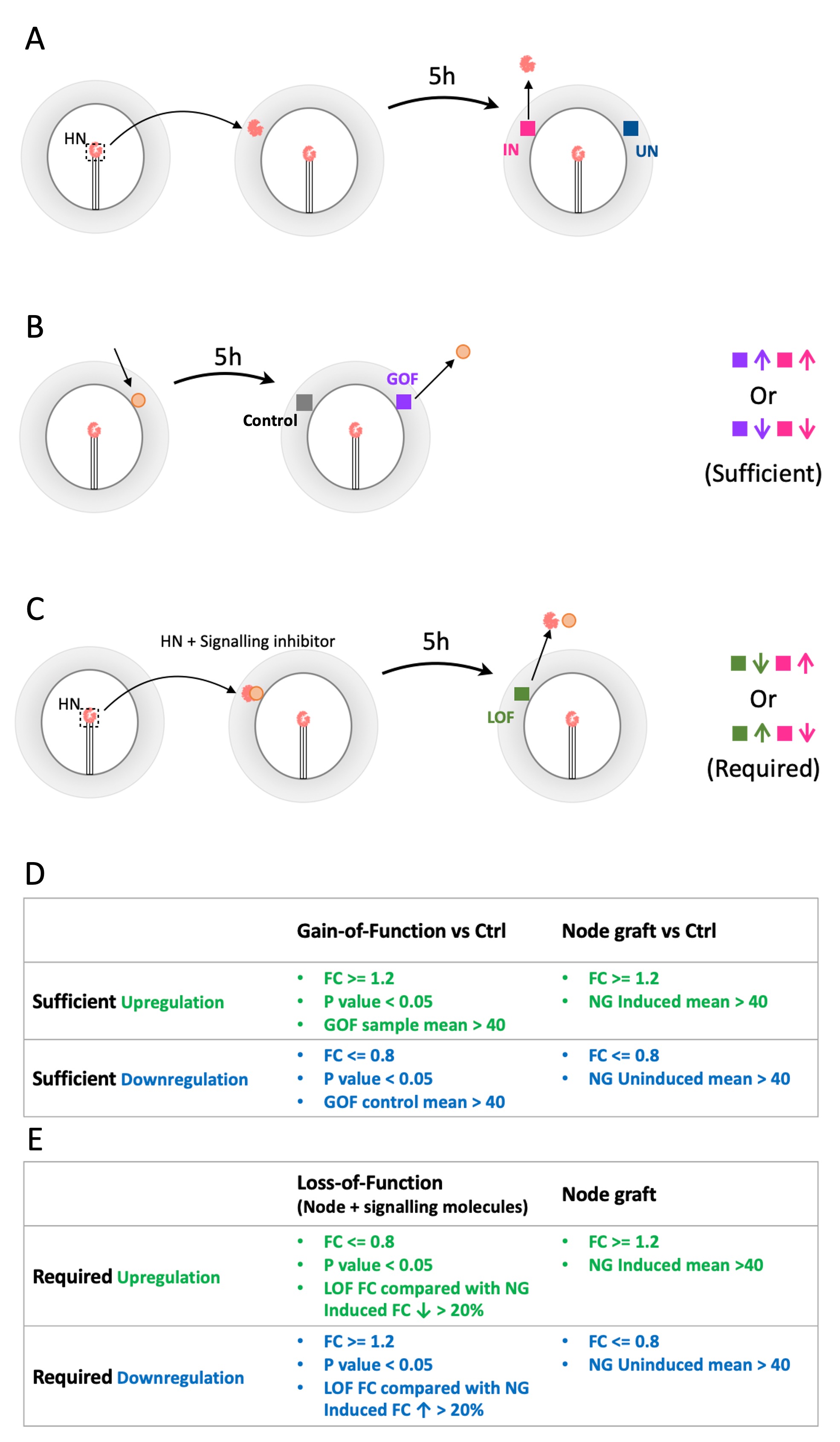

### Supplementary Figure S5

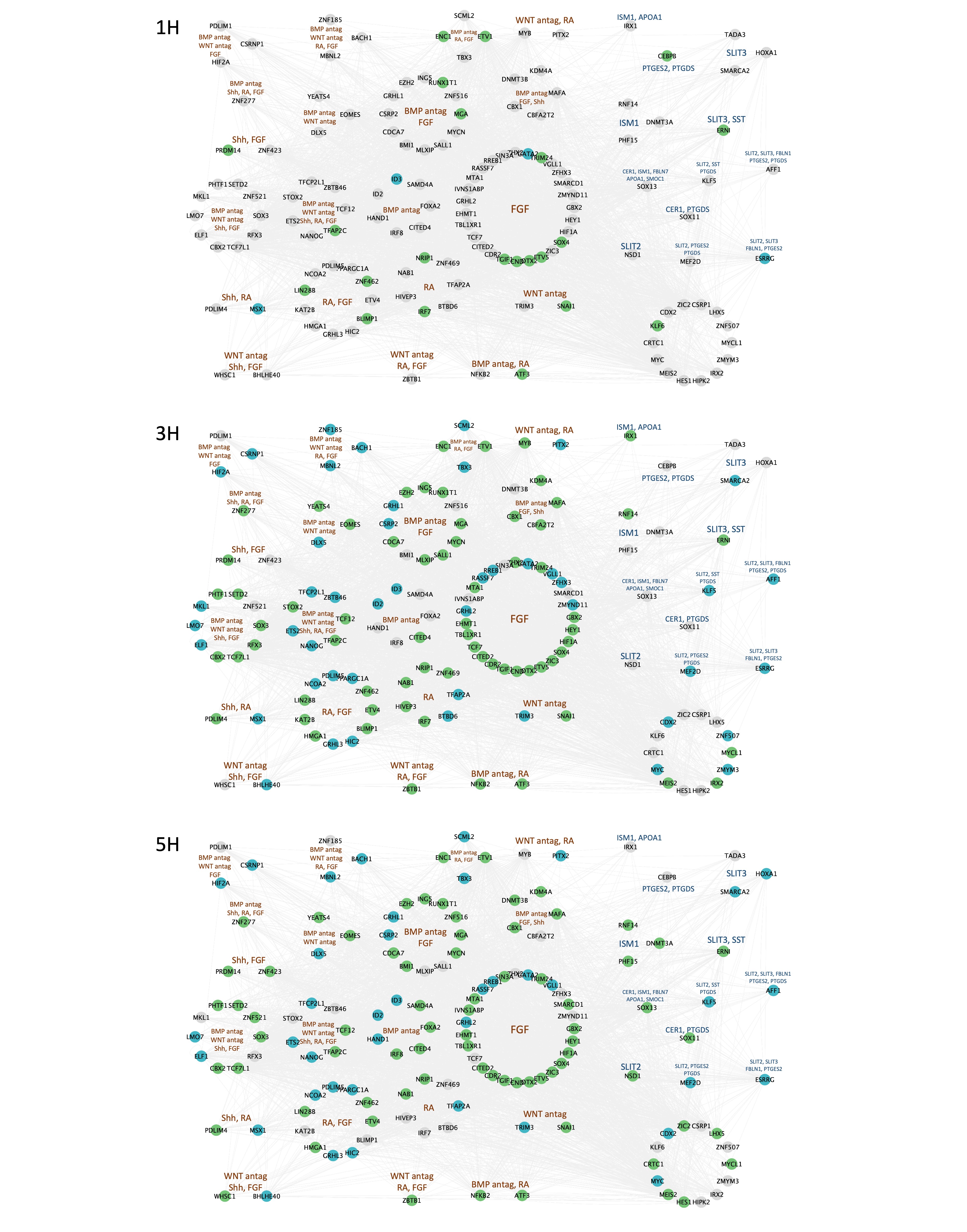

### Supplementary Figure S6

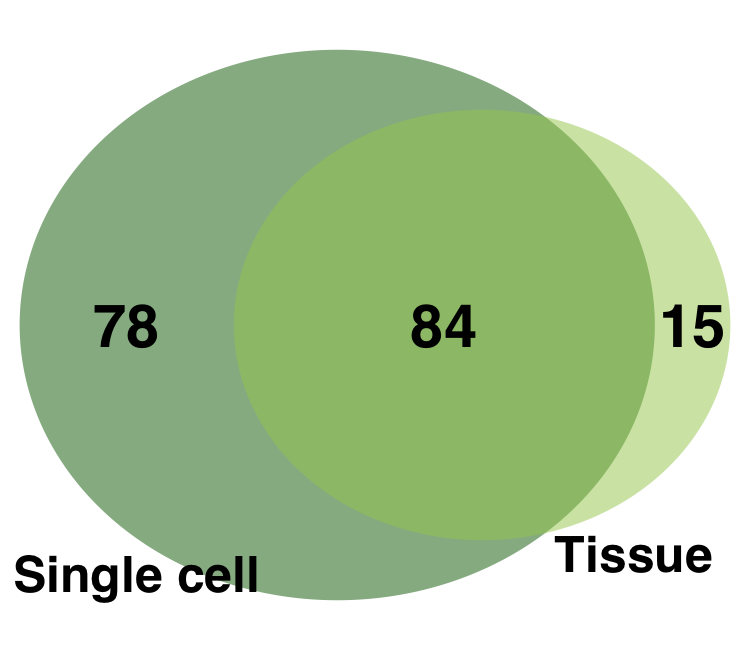
